# From Weak Interactions to Strong Affinity: Deciphering the Streptavidin-Biotin Interaction through NMR and Computational Analysis

**DOI:** 10.1101/2024.12.19.629369

**Authors:** Aleksandra L. Ptaszek, Sarah Kratzwald, Filip Sagan, Mario Migotti, Pedro A. Sánchez-Murcia, Robert Konrat, Gerald Platzer

## Abstract

Understanding weak interactions in protein-ligand complexes is essential for advancing drug design. Here, we combine experimental and quantum mechanical approaches to study the streptavidin-biotin complex, one of the strongest known protein-ligand binders. Using a monomeric streptavidin mutant, we analyze ^1^H NMR chemical shift perturbations (CSPs) of biotin upon binding, identifying unprecedented upfield shifts of up to -3.2 ppm. Quantum chemical calculations attribute these shifts primarily to aromatic ring currents, with additional contributions from charge transfer effects linked to weak interactions. The agreement between experimental and computed chemical shifts validated the X-ray structure as a reliable basis for detailed computational analyses. Energy decomposition analysis reveals that electrostatics dominate the biotin-streptavidin interaction, complemented by significant orbital and dispersion contributions. Notably, weak non-covalent interactions—such as CH· · · S, CH· · · *π*, and CH· · · HC contacts—driven by London dispersion forces, contribute ∼44% to the complex’s stability.

## Introduction

Nuclear Magnetic Resonance (NMR) spectroscopy is an indispensable tool in chemistry and structural biology, providing atomic-level insights into molecular dynamics, conformational changes, and intricate intermolecular interactions. Its unique ability to examine molecules under near-physiological conditions makes NMR particularly valuable for studying complex systems such as proteins and protein-ligand interactions.^1^

Traditionally, scalar couplings (J-couplings) have been extensively employed to probe bonding relationships and conformational states, particularly in small molecules. The presence of scalar couplings, which were initially observed in covalent bonds, is now known to extend to hydrogen bonds, demonstrating their complex nature driven by both electrostatic forces and electronic density transfer.^2^ Over recent years, *through-space* scalar couplings have been identified in interactions at the very edge of hydrogen bonding, including CH· · · *π* contacts, where a soft acidic CH bond interacts with a soft basic *π*-system.^3,4^ Even more intriguingly, such couplings have been detected between homopolar CH· · · HC contacts, suggesting that these interactions, though weak, exhibit characteristics of true bonding.^5–8^ While J-couplings provide rich structural information, their application is challenging in large protein systems. The spectral crowding and faster relaxation times in these systems reduce the practical utility of J-couplings, particularly for routine applications like drug discovery.

Another valuable NMR observable that provides extensive insight into the local chemical environment is the chemical shift. For example, a proton chemical shift serves as an effective indicator of hydrogen bonding, where stronger hydrogen bonds lead to increased deshielding effects. Additionally, proximity of aromatic rings induces shielding effects due to ring currents, perturbing chemical shifts.^9^ Upon protein binding, ligand proton chemical shifts can change due to a combination of factors, including loss of hydrogen bonds with solvent and formation of new hydrogen bonds within the binding pocket. Moreover, binding modes frequently involve aromatic rings, leading to additional shielding effects from ring currents. These combined factors provide a rich source of structural information that can be harnessed to study protein-ligand interactions in detail.

In this study, we explore the interaction between streptavidin and biotin, which is recognized as one of the most robust non-covalent interactions found in nature, and has been a target of numerous studies in the past Figure 1.^10–14^ The high affinity of this interaction is exploited within a wide range of biotechnological applications (e.g. *Strep-tags*).^15^ Despite long-lasting scientific interest in the streptavidin-biotin complex, the origin of the high binding strength remains elusive. Besides a high shape complementarity, a large number of classical hydrogen bonds involving the biotin head-piece nitrogens and oxygen as well as the carboxy terminus of the aliphatic tail significantly contribute to the high binding enthalpy of the system. It is the "hydrophobic tail" however, that adds several non-classical hydrogen bonding interactions that are rarely discussed in a protein-ligand binding context which have been shown to majorly contribute to the total free binding energy of the system. The term "hydrophobic interaction" is often misused in the context of CH, CH_2_ and CH_3_ groups interacting with their environment. In addition to hydrogen bonds to classical acceptors (O,N,S) and the distinct case of Fluorine, ^16,17^ these groups also form CH· · · *π* ^3,9^ interactions with nearby aromatic and homopolar CH· · · HC interactions^18–20^ to nearby aliphatic residues. Interactions of carbon-bound protons are of special interest given their high frequency in protein-ligand complexes accounting for over 50% of all non-covalent interactions present in guest-host systems. ^21^ In addition, their presence can be detected straightforwardly via simple 2D NMR experiments.^9^ Since native Streptavidin (PDB ID: 3RY2) forms a tetramer in solution, we opted to use a slightly modified, monomeric form of Streptavidin (PDB ID: 4JNJ) to make the protein construct amenable for NMR measurements. ^22,23^ In the streptavidin-biotin complex under study (PDB ID: 4JNJ), only two out of fifteen hydrogen atoms form classical NH· · · O hydrogen bonds with Asn45 (Ser45 in PDB ID: 3RY2) and Asp128, Figure 1 c). Beyond that, the X-ray structure reveals weaker CH· · · O interactions with Thr48 (Val47 in PDB ID: 3RY2) and Gly49, and a CH· · · S contact with Cys59, Figure 1 c,d). Further, biotin’s binding pocket showcases multiple CH· · · *π* interactions with aromatic rings of Trp79 and Trp108 residues. Additionally, homopolar CH· · · HC interactions with Leu110 highlight the complex’s reliance on non-classical, dispersion-driven forces that collectively support biotin’s snug fit within streptavidin. Amid the ongoing scientific debate, considerable research has been devoted to deciphering the nature of CH· · · HC interactions. Until recently, they were mostly perceived with the notion of "steric crowding", but more and more works emphasize the stabilization provided by such contacts, stemming from London dispersion, and also from electronic interactions.^24–38^

**Figure 1:**
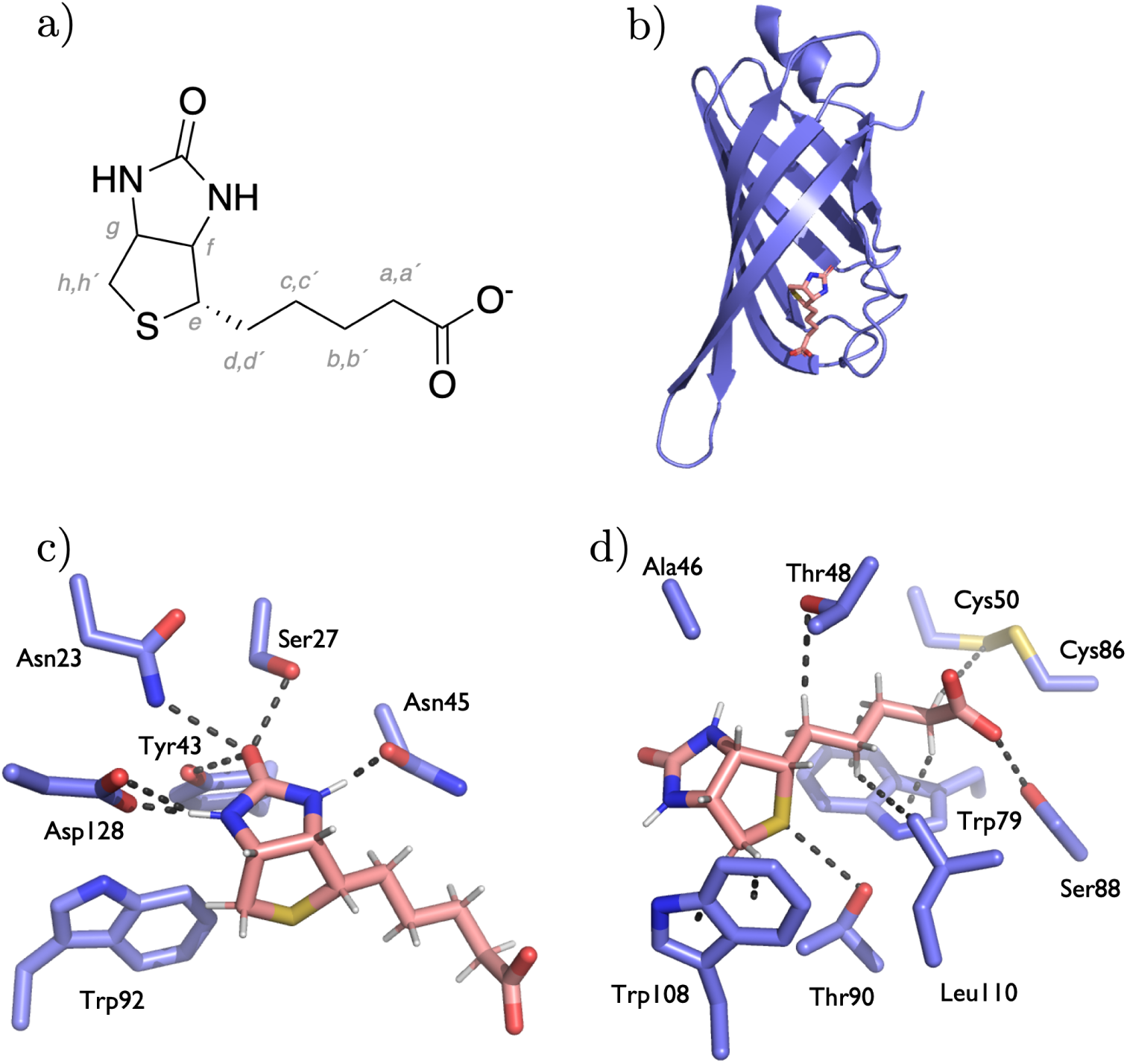
(a) Chemical structure of biotin at pH 7.4 with H-labeling used throughout this work; (b) X-ray structure of mSA2 and biotin (PDB ID 4JNJ);^22^ (c) Interactions of biotin’s ureido ring with streptavidin pocket; (d) Interactions of biotińs tetrahydrothiophene ring and valeric acid side chain with surrounding streptavidin residues.

Previously, we reported the PI by NMR methodology which allows the quantification of the strength of these CH· · · *π* interactions.^9^ Recently, the approach has been developed further. Thereby, protein deuteration and ligand-based ^1^H NMR experiments were used to probe CH*^ligand^*-*π^protein^* interactions by ^1^H NMR experiments.^39^ We showed that ligand proton chemical shifts, when combined with computational shift predictions and molecular dynamics simulations, can refine protein-ligand interfaces, providing valuable insights into solution structures. This study employs a ^1^H-^1^H NOESY experiment to quantify chemical shift perturbation (CSP) for the carbon-bound protons of biotin in both its free and bound forms. To validate the solution structure against crystallographic data, we compare the experimentally obtained shifts with quantum mechanically (QM) predicted shifts based on the X-ray structure of the streptavidin-biotin complex. Following this validation, we use modern chemical bonding descriptors such as ETS-NOCV^40^ and LED-DLPNO-CCSD(T) ^41–48^ to explore the interactions of biotin with the binding pocket, focusing on the non-classical hydrogen bonds. We also decompose the carbon-bound proton chemical shifts to separate the shielding effects of environmental ring currents from deshielding due to hydrogen bond formation. Our findings show that the large CSPs are primarily due to ring current effects, with hydrogen bonding playing a secondary role. In conclusion, by integrating NMR and computational methods, we offer a comprehensive understanding of the forces that govern biotin–streptavidin molecular interactions.

## Results and Discussion

### Detection of Chemical Shift Perturbations of biotin’s H-Atoms

In this work we study a monomeric mutant of the streptavidin protein (mSA2)^22,23^ which, due to its more favorable relaxation properties is more suitable for NMR applications compared to the homotetrameric wild-type (WT). In the tetrameric WT streptavidin, a tryptophan residue (Trp120) of an adjacent subunit forms additional stacking interactions with biotin which is associated with the extraordinarily high wild-type affinity towards biotin. ^49^ The mSA2 mutant used in this study has a binding affinity in the low nanomolar range (K*_D_* = 1.9 ± 0.4 × M*^−^*^9^),^23^ whereas the WT streptavidin has a binding affinity in the picomolar range (K*_D_* = 4 × 10*^−^*^14^ M).^50^ The mSA2 mutant is a hybrid of rhizavidin and streptavidin (Figure S1). Unlike streptavidin, rhizavidin binds to biotin with a single subunit. In creating mSA2, amino acids in the biotin binding pocket of streptavidin were replaced with those from rhizavidin. Notably, many of these residues are common to both proteins (see Figure S2).^51^

To determine chemical shift perturbations (CSP) of biotin upon binding to streptavidin, we perdeuterated mSA2 to render all protein ^1^H signals NMR inactive. To evaluate CSPs for each of biotin’s non-exchanging hydrogen atoms, ^1^H peak assignment of biotin’s protons was performed as described previously^39^ via a ^1^H-^1^H NOESY experiment of a perdeuterated 1:2 streptavidin-biotin complex. This allows for an unambiguous correlation of biotin’s unbound H-signals with biotin’s H-signals bound to streptavidin (see Figure S4).

Figure 2 shows the ^1^H NMR spectra of free biotin (top) and biotin bound to deuterated streptavidin (bottom). Remarkably, CSPs of -3.2, -2.9, -2.1 and -1.9 ppm for biotin’s H-atoms *h, b, h′* and *a* could be observed. Large ^1^H upfield shifts indicate shielding for affected Hatoms which is induced by the *π* cloud of aromatic ring systems when located above/below of the respective H-atom thereby forming favorable CH· · · *π* interactions with streptavidin’s aromatic indole moieties of Trp79 (for *a* and *b*) and Trp108 (for *h* and *h′*) respectively (see Figure 2 b). The interaction can be considered as a special case of hydrogen bonding where the C-H group serves as the H-bond donor and the aromatic ring as the H-bond acceptor. ^52^

**Figure 2:**
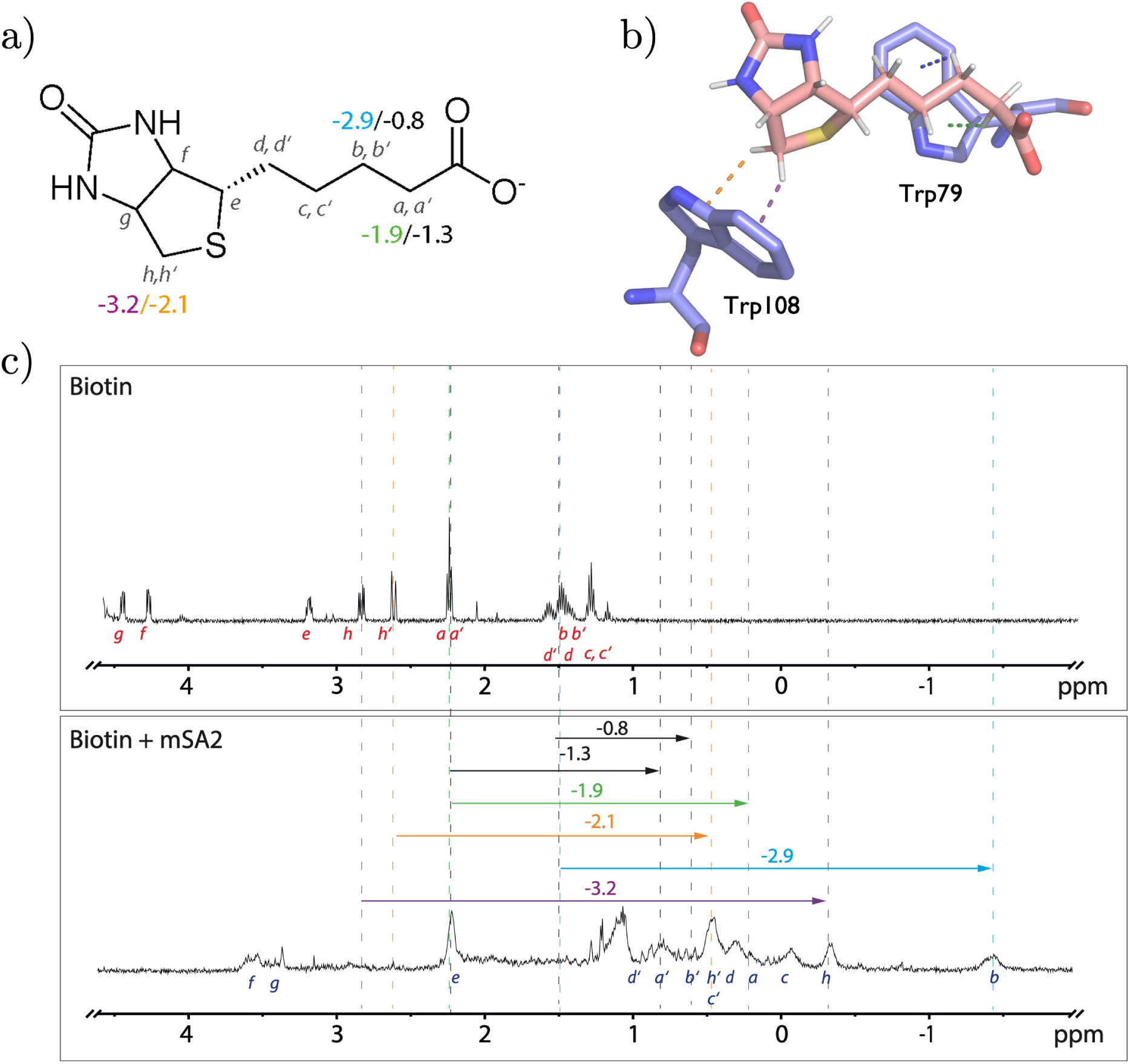
(a) Chemical structure of biotin highlighting the protons undergoing the largest CSP. (b) Crystal structure of biotin-mSA2. Protons a, b, h and h’ are placed right in top of the indole moiety of Trp79 (proton a and b) and Trp108 (proton h and h’). (c) *top:* ^1^H-NMR of biotin in PBS buffer, *bottom:* ^1^H-NMR of biotin bound to deuterated mSA2. The dashed lines depict the ^1^H CSPs of biotin’s free to bound state for proton a, b, h and h’. Detailed results provided in Supplementary Table S1.

All H-atoms of biotin’s aliphatic tail (*a, a′, b, b′, c, c′, d, d′ and e*) undergo an upfield shift upon binding (min - 0.49 ppm; max - 2.89 ppm). The general observation that carbon-bound ligand protons shift to lower ppm values upon binding can be attributed to two factors; 1) Protons moving from the solvated free state to a solvent-inaccessible bound state lose hydrogen-bonding to water resulting in an apparent upfield shift (hydrogen-bonding itself has a deshielding character) and 2) ring systems exert their effect up to a distance of 4Å away and can influence protons that are not directly interacting with them. (*b′,c, c′, d, d′, e*). The last section of the results discusses the detailed decomposition of the ^1^H chemical shifts of biotin in the bound form into environmental shielding and deshielding due to the hydrogen bond.

### ^1^H NMR Chemical Shifts Prediction

In a previous study, we investigated the ^1^H chemical shifts (CSs) of the BI-9321 molecule bound to the PWWP1 domain of NSD3.^39^ By validating calculated CSs against experimental ligand ^1^H CSs in the bound form, we identified significant discrepancies for certain protons. To address these, we applied a quantum mechanics/molecular mechanics (QM/MM) molecular dynamics (MD) ensemble approach to refine the initial X-ray structure, which significantly improved agreement with the experimental ^1^H CSs in solution, achieving an absolute error (RMSE) of less than 0.5 ppm for each proton and a root mean square error (RMSE) of 0.28 ppm.

In this study, we used the same theoretical approach to compute chemical shieldings for the biotin-mSA2 X-ray structure (PDB ID: 4JNJ)^22^ and validated the results with experimental ^1^H CSs. Figure 3 a) demonstrates a strong correlation between calculated and experimental values, with an RMSE of 0.24 ppm. Furthermore, absolute errors for all protons remained below 0.5 ppm (Figure 3 b). These results confirm that the X-ray structure of biotin-mSA2 provides an accurate representation of the complex in solution.

**Figure 3:**
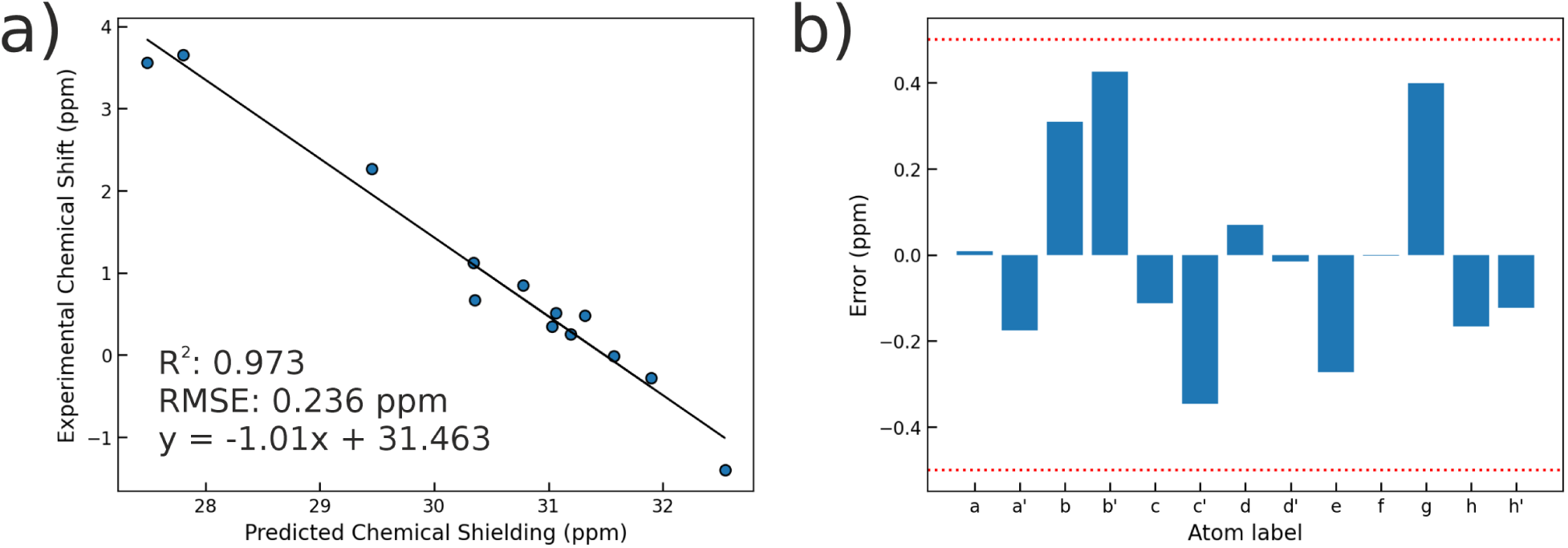
a) Correlation between experimental NMR ^1^*H* CSs (ppm) and calculated chemical shielding (ppm) from the X-ray crystal structure (PDB ID: 4JNJ). b) Differences between calculated and experimental chemical shifts (ppm). GIAO/wB97X-D/def2-TZVP level of theory was applied. ^53–58^

### Biotin-mSA2 Interaction Energy Analysis

To gain a comprehensive understanding for the physico-chemical properties driving Biotin-mSA2 complex formation, we used the ETS-NOCV charge and energy decomposition scheme ^40^ at the DFT level as well as the LED energy decomposition based on the DLPNO approximation of the *gold-standard* CCSD(T) method.

In the first step, we consider the interaction of the biotin molecule (*fragment 1*) with the binding site and water molecules in direct contact with biotin (*fragment 2*) using the geometric cluster previously defined for the ^1^H CS calculations, as described in the Methods section, Figure 4 a). The overall interaction energy reaches -86.67 kcal/mol and is dominated by electrostatic interactions (40%) followed by orbital (36%) and dispersion (24%) components, Table 1 a). The deformation density reveals numerous charge-flow channels associated with conventional *N/O* − *H* · · · *O* hydrogen bond formation in the ureido ring and carboxylate regions, as well as weaker *C* − *H* · · · *O* and *O* − *H* · · · *S* hydrogen bonds, Figure 4 a). In order to exclude the strong hydrogen bonds with water molecules from the interaction energy, we include water molecules in *fragment 1* getting the total interaction energy equal to -83.36 kcal/mol (Table 1 b), Figure 4 b)) which is in an excellent agreement with the *state-of-the-art* DLPNO-CCSD(T) interaction energy equal to -83.23 kcal/mol. In comparison to the initial system (Table 1 a)), the absence of two prominent hydrogen bond donors results in a rise of approximately 20 kcal/mol in both electrostatic and orbital energy terms, which is counterbalanced by the reduction in Pauli repulsion.

**Figure 4:**
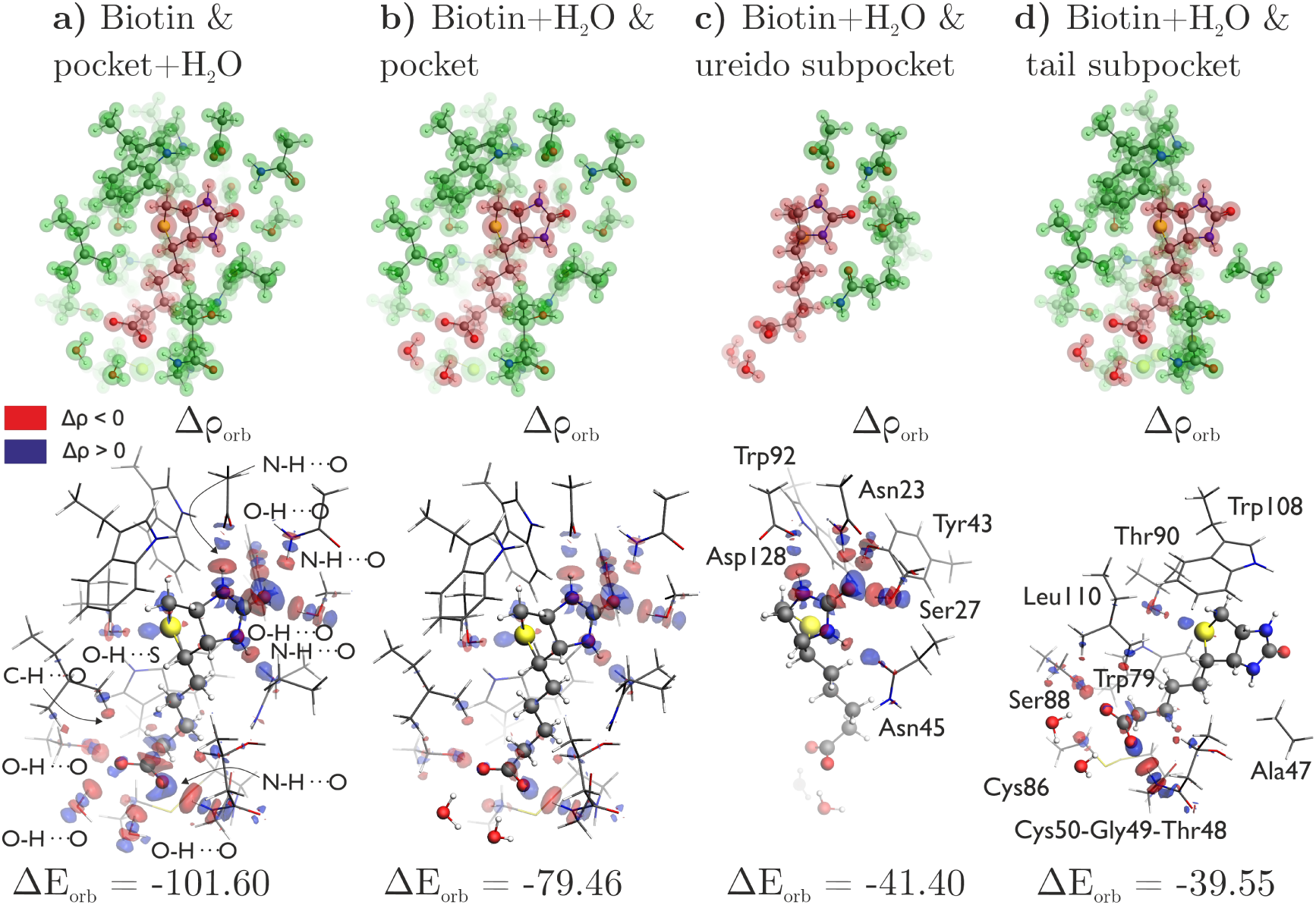
The initial row illustrates the ETS-NOCV fragmentation using red and blue, as described in Table 2. Below, the overall deformation density Δ*ρ_orb_* contours with the corresponding Δ*E_orb_* for 4 models. The biotin molecule is represented by balls and sticks, and the binding site is represented by sticks. The inflow of electronic density in blue and the outflow in red. Energy in kcal/mol.

**Table 1:** ETS energy decomposition results describing the interaction of biotin with the binding site: a) biotin molecule interacting with two water molecules and binding pocket, b) biotin and two water molecules interacting with the binding pocket, c) biotin and two water molecules interacting with the ureido subpocket (Asn23, Ser27, Tyr43, Asn45, Trp92, Asp128), d) biotin and two water molecules interacting with the tail subpocket (Ala47, Thr48, Gly49, Cys50, Cys86, Trp79, Ser88, Thr90, Trp108, Leu110). The percentages represent the proportionate contribution to the overall attractive interactions (*E_elstat_* + Δ*E_orb_* + Δ*E_disp_*). All energies in kcal/mol.

| Energy term | a) Biotin ...<br>pocket+H<sub>2</sub>O | b) Biotin+H<sub>2</sub>O ...<br>... pocket | c) Biotin+H<sub>2</sub>O ...<br>ureido subpocket | d) Biotin+H<sub>2</sub>O ...<br>tail subpocket |
| --- | --- | --- | --- | --- |
| $\Delta E_{total}$ | -86.67 | -83.36 | -23.49 | -61.27 |
| $\Delta E_{elstat}$ | -113.16 (40%) | -89.17 (37%) | -34.85 (36%) | -55.57 (38%) |
| $\Delta E_{orb}$ | -101.60 (36%) | -79.46 (33%) | -41.40 (43%) | -39.55 (27%) |
| $\Delta E_{disp}$ | -68.72 (24%) | -71.93 (30%) | -20.35 (21%) | -51.59 (35%) |
| $\Delta E_{Pauli}$ | 196.82 | 157.20 | 73.11 | 85.45 |

**Table 2:** ETS energy decomposition results describing the interaction of biotin (**fragment 1**) with individual residues of the tail subpocket. Within parentheses, the contribution to the overall biotin+*H*_2_*O* · · · pocket stability is outlined. Energies in kcal/mol.

| **Interacting**<br>**fragment 2** | $\Delta E_{total}$ | $\Delta E_{elstat}$ | $\Delta E_{orb}$ | $\Delta E_{disp}$ | $\Delta E_{Pauli}$ |
| --- | --- | --- | --- | --- | --- |
| Gly49 | -14.33 (16%) | -21.61 | -14.12 | -6.84 | 28.24 |
| Ser88 | -11.58 (13%) | -9.11 | -5.34 | -2.87 | 5.73 |
| Cys50-Cys86 | -8.16 (9%) | -5.18 | -4.41 | -7.30 | 8.73 |
| Trp79 | -7.44 (9%) | -4.71 | -3.14 | -10.54 | 10.95 |
| Thr48 | -6.62 (8%) | -3.74 | -3.15 | -5.13 | 5.40 |
| Trp108 | -5.47 (6%) | -2.99 | -1.70 | -7.38 | 6.59 |
| Leu110 | -5.07 (6%) | -2.61 | -2.63 | -5.33 | 5.50 |
| Thr90 | -3.96 (5%) | -5.66 | -5.54 | -4.23 | 11.48 |
| Ala47 | -1.03 (1%) | -1.80 | -0.84 | -3.10 | 4.71 |

Delving deeper into the contributions of individual regions within the binding pocket to the overall stability, we analyse two distinct cases: (1) the interactions of biotin with only those residues from the binding pocket that interact with its ureido ring, referred to in the text as the *ureido subpocket* (Figure 1 c)), and (2) interactions of biotin with the remaining residues of the binding pocket, referred to as the *tail subpocket* (Figure 1 d)). Total interactions between the biotin and the ureido subpocket amount to approximately 28% of biotin-mSA2 stability, Table 1 c). Analysis of the deformation density uncovers charge-flow channels related to the formation of five classical hydrogen bonds (Figure 4 c)), yielding an energy of -41.40 kcal/mol, which accounts for approximately 43% of the overall attractive forces in this system. Additionally, electrostatic interactions contribute around 36% of the overall attraction. The dominance of orbital and electrostatic factors aligns with prior analyses of classical hydrogen bonds. ^59^

Focusing now on the interactions of biotin with the tail subpocket, electrostatic forces emerge as the primary contributor, constituting 38% of the interactions, likely influenced by the negatively charged carboxylic acid moiety at physiological pH (pK*_a_* ∼4.5) (Table 1 d). Interestingly, calculations indicate that approximately 35% of the stabilizing interactions are attributed to dispersion forces. To validate this empirical estimation of London dispersion, we cross-referenced it with results obtained using the LED-DLPNO-CCSD(T) level of theory, and found a slight overestimation on the part of DFT (-51.6 kcal/mol vs -42.8 kcal/mol). Nevertheless, the dispersion contribution within the biotin - tail subpocket model remains twice as high as for the biotin - ureido subpocket, owing to much higher degree of packing around the hydrophobic tail. As evidenced by the Dispersion Interaction Density (Figure S6 b), the entirety of tail section is engaged in dispersive stabilization, whereas in the ureido subpocket case, only hydrogen bond donors and acceptors are stabilized in this way (Figure S6 a). Furthermore, orbital part of the energy is on par with the ureido subpocket fragment, and constitues 27% of total stabilization within the tail subsystem (Table 1 d), Figure 4 d). Figure S5 illustrates the NOCV-based deformation density channels, highlighting both *σ* and notably weaker (∼-1 kcal/mol) *π* components of the *N* − *H* · · · *O* hydrogen bonds. Additionally, the analysis reveals the involvement of sulfur in hydrogen bonding interactions, not only in the *O* − *H* · · · *S* bond with Thr90, but also in weaker (∼ -1.6 kcal/mol) *C* − *H* · · · *S* bonds with Trp79 and the methyl group of Thr90. Moreover, the NOCV channels corresponding to the *C* − *H* · · · *π* interactions are identified, Figure S5.

We also analysed how biotin interacts with individual residues from the tail subpocket to elucidate their role in overall stability, Table 2, Figure S7. The results show that fragments interacting with biotin mainly through *N/O* − *H* · · · *O* classical hydrogen bonds, such as Gly49 and Ser88, are predominantly governed by electrostatic interactions. Collectively, these fragments provide a stabilization comparable to 29% of the mSA2 · · · biotin stability. Deformation density analysis additionally reveals charge transfer channels associated with the weaker *C* −*H* · · · *O/S* hydrogen bonds with Thr48 and Cys50-Cys86, Figure S7. The *O* − *H* · · · *S* type hydrogen bond between Thr90 and biotin is equally dominated by electrostatic and orbital components and compares to 5% of the global binding. The residues Trp79, Trp108, and Ala47 compare to 15% of the total binding energy, predominantly contributing through *C* −*H* · · · *π* contacts. An additional 6% of the equivalent of binding energy is derived from *C* − *H* · · · *H* − *C* interactions with Leu110. Both of these interaction types are primarily governed by London dispersion forces.

### Probing Hydrogen Bonding Effects via ^1^H NMR Shifts

In this paragraph, we aim to elucidate the physical factors behind the chemical shifts observed in biotin carbon-bound protons upon protein binding. By following the approach proposed by Scheiner,^60^ we identify two primary components affecting the chemical shift of interest: 1) an environmental shielding effect and 2) deshielding due to hydrogen bond formation (details are given in the Methods section), Table 3.

**Table 3:** Decomposition of the factors contributing to chemical shifts of individual biotin C-bounded protons along with the lists of their H-bond acceptors and hydrogen bond lengths. In the case of C-H· · · *π* interactions, the distance from the proton to the center of the ring is considered.

| Proton label | Environmental shielding effect | Deshielding due to H-bond formation | Exp. CSP | H-bond acceptor | H-bond length (Å) |
| --- | --- | --- | --- | --- | --- |
| a | -1.93 | 0.64 | -1.88 | $\pi$(Trp79) | 2.9 |
|  |  |  |  | O(Ser88) | 3.0 |
| a' | -0.93 | 0.61 | -1.28 | S(Cys50) | 2.9 |
|  |  |  |  | H-C(Cys50) | 2.2 |
| | | | | $\pi$(Trp79) | 4.1 |
| b | -2.32 | 0.52 | -2.89 | $\pi$(Trp79) | 3.1 |
|  |  |  |  | N(Asn45) | 2.9 |
| b' | -0.82 | 0.49 | -0.82 | O(Gly49) | 2.4 |
|  |  |  |  | N(Asn45) | 2.8 |
| | | | | $\pi$(Trp79) | 4.8 |
| c | -1.42 | 0.46 | -1.31 | $\pi$(Trp79) | 3.1 |
| c' | -0.89 | -0.27 | -0.82 | H-C(Leu110) | 2.3 |
|  |  |  |  | O(biotin CO<sub>2</sub><sup>-</sup>) | 3.2 |
| | | | | $\pi$(Trp79) | 4.6 |
| d | -0.84 | 0.62 | -1.17 | $\pi$(Trp79) | 3.9 |
|  |  |  |  | O(Asn45) | 2.9 |
| d' | -0.32 | 0.38 | -0.49 | O(Thr48) | 2.7 |
|  |  |  |  | N(Asn45) | 3.0 |
| | | | | $\pi$(Trp79) | 5.3 |
| e | -0.72 | 0.06 | -0.98 | $\pi$(Trp108) | 4.9 |
|  |  |  |  | H-C(Leu110) | 2.6 |
| f | -0.31 | 0.12 | -0.70 | $\pi$(Trp108) | 5.4 |
|  |  |  |  | H-C(Thr48) | 2.5 |
| g | -0.63 | 0.28 | -0.94 | $\pi$(Trp108) | 3.6 |
| h | -3.35 | 0.25 | -3.17 | $\pi$(Trp108) | 2.7 |
| h' | -1.99 | 0.66 | -2.15 | $\pi$(Trp108) | 3.0 |

The environmental shielding component accounts for the magnetic shielding effects arising from the surrounding molecular environment. These include ring current effects caused by aromatic residues, as well as positional shielding from nearby atoms or groups. Computationally, this effect was isolated by replacing the biotin protons with dummy atoms while retaining the protein binding pocket. The highest shielding values were observed for protons **h**, **b**, **h’**, and **a**, which explains the significant experimental CSPs noted in these regions, Table 3. The shielding effects extend beyond direct hydrogen bond lengths, demonstrating the long-range influence of aromatic rings, with observed values ≥ 0.3, even for protons significantly distant from the aromatic ring. These findings align with our empirical estimate of the ring current effect as predicted by the classical Pople model (Table S3).

The deshielding component arises from hydrogen bond formation between biotin protons and hydrogen bond acceptors within the protein binding pocket. This interaction introduces localized charge transfer and polarization effects, leading to reduced electron density around the protons and thus a deshielding effect. The analysis of various types of hydrogen bonds presented in Schreiner’s work^60^ suggests that the “deshielding due to H-bond” correlates with the bonding strength. The largest "deshielding effect due to the H-bond" is observed for protons **a** and **h‘**, both of which are located directly above the pyrrole rings of tryptophan amino acids (Table 3, Figure 5). In the paper of Scheiner,^60^ this position was identified as the most energetically favourable for the water · · · indole dimer. Moreover, the hydrogen bond character of protons **a**, **b**, and **d** is further enhanced by weak CH · · · O and CH · · · N bonds, as evidenced by the deformation density channels visualized in Figure 5. Interestingly, the lowest, but nonzero deshielding due to H-bond formation is identified for protons **f** and **e** that exhibit only homopolar CH· · · HC interactions, corresponding charge-flow channels are presented in Figure 5.

**Figure 5:**
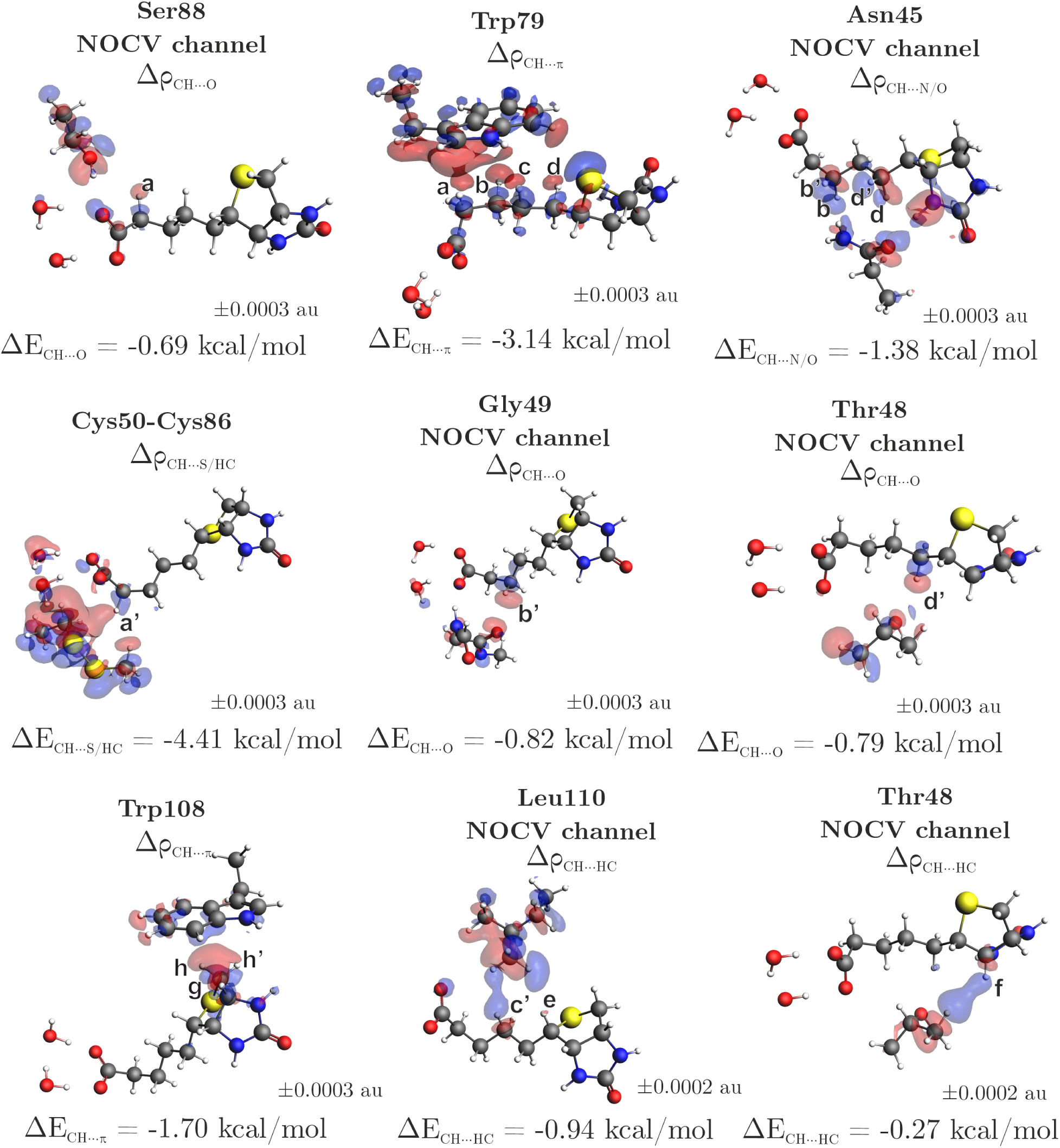
The contours of deformation density channels revealing the charge transfer of carbon-bounded biotin protons caused by hydrogen bonds formation. The inflow of electronic density is shown in blue and the outflow in red. Carbon-bounded protons of interest are labeled.

Surprisingly, the proton **c’** does not follow the expected trend (positive sign) for "deshielding due to H-bond formation", Table 3. For this specific proton, the deshielding effect due to hydrogen bonding in the bound state is masked by a stronger deshielding effect in the unbound state. In the absence of a binding pocket, the carboxylate group of biotin carries a negative charge, causing it to polarize the proton **c’** leading to the formation of an intramolecular hydrogen bond in the free form. In the bound form, this interaction is relieved and the **c’** proton interaction is replaced with weaker H-bond acceptors in the binding pocket. This leads to an apparent shielding as reflected by the negative value (-0.27) for proton **c’** in Column 3 of Table 3. This phenomenon is observed in the chemical shift calculation of the free form because in our model the ligand is maintained in the same conformation as in the bound form, thus preventing the rotation of the biotin tail.

## Conclusions

In this study, we aimed to provide a detailed understanding of the molecular interactions governing the binding of biotin to the monomeric streptavidin mutant (mSA2), a system known for its exceptionally strong non-covalent protein-ligand interaction. Through ^1^H NMR chemical shift analysis and quantum chemical calculations, we uncovered important insights into the nature of these interactions.

Upon binding to mSA2, all carbon-bound protons in biotin shifted to significantly lower ppm values. Chemical shifts were accurately predicted using quantum chemical methods based on the X-ray structure. The strong agreement between the predicted and experimental shifts confirmed that the X-ray structure is a reliable representation of the streptavidin-biotin complex in solution, allowing for further computational studies of the binding mechanism.

The decomposition of proton chemical shifts in the bound form revealed that the strongest chemical shift perturbations are primarily due to the ring current effect from nearby aromatic residues. Moreover, the chemical shifts of all carbon-bound protons participating in weak interactions, like CH· · · O, CH· · · S, CH· · · *π*, and even homopolar CH· · · HC, are impacted by the contributions arising from charge transfer as well.

Energy decomposition analysis revealed that the biotin-mSA2 interaction is primarily driven by electrostatic forces, with additional contributions from orbital and dispersion interactions. These findings were corroborated by LED-DLPNO-CCSD(T) calculations, which showed strong agreement with the density functional theory results. Further NOCV analysis and fragmentation of the system allowed us to pinpoint weaker non-covalent interactions, such as CH· · · S, CH· · · *π*, and homopolar CH· · · HC, alongside classical hydrogen bonds. Our analysis quantified that non-classical hydrogen bonds, including OH· · · S, CH· · · S, CH· · · *π*, and CH· · · HC, contribute approximately 44% to the overall stability of the complex. We demonstrated that these interactions are predominantly governed by London dispersion forces, with smaller contributions from electrostatics and orbital interactions. This contrasts with classical hydrogen bonds, which typically rely on electrostatic and orbital components.

These findings highlight the nuanced role of weak non-covalent interactions in stabilizing protein-ligand complexes. Such a detailed understanding of binding mechanisms and the nature of non-covalent interactions is crucial for drug discovery, as it enables the design of molecules that leverage these weak forces to achieve high-affinity binding.

## Materials and Methods

### Plasmid Design

DNA sequences encoding the fusion construct H6-MBP-3C-mSA2 were synthesized by GeneScript and cloned into the pETM44 vector via Sequence and Ligation Independent Cloning (SLIC). This construct integrates a hexahistidine (H6) tag and maltose-binding protein (MBP) for purification, linked to the mSA2 protein through a 3C protease cleavage site to allow for H6-MBP post-purification. The integrity and sequence accuracy of the insert were confirmed by sequencing.

### Protein Expression and Purification

Recombinant H6-MBP-mSA2 (monomeric streptavidin 2) was expressed in *E. coli BL21 (DE3)* and contains an N-terminal 3C protease cleavable H6-MBP tags. ^15^N labeled H6-MBP-3C-mSA2 was expressed in M9 medium containing ^15^NH_4_Cl (1 g/l). Cells were grown to an OD_600_ (Optical Density at 600 nm) of 0.9 and induced with 0.4 mM IPTG (isopropyl-*β*-D-1-thiogalactopyranosid). The expression medium was additionally supplemented with biotin (1 mg/l) for the mSA2-biotin construct. Cells were harvested after 16 h of expression at 20°C. Perdeuterated H6-MBP-3C-mSA2 was expressed by transferring 100 *µ*l of transformed *E. coli BL21(DE3)* cells from M9 medium (no isotope-enriched material, 100% H_2_O) to M9^50%^*^D^*^2^*^O^* (no isotope-enriched material, 50% D_2_O, 50% H_2_O) until an OD_600_ of 0.7 was reached. A small amount of *E. coli* cells were transferred to 50 ml M9^100%^*^D^*^2^*^O^* (no isotope-enriched material, 100% D_2_O) preculture. The next day, a small aliquot of the cells was pelleted and transferred to the M9^100%^*^D^*^2^*^O^* expression medium (D-glucose-d7, 100% D_2_O). After reaching an OD_600_ of 0.9, induction was performed using 0.4 mM IPTG. Cells were harvested after 20 h at 20°C, and the pellet was resuspended in 30 ml of lysis buffer (20 mM TrisHCl pH 7.4, 300 mM NaCl, 10 mM Imidazole, 1 mM TCEP, 0.5% (v/v) CHAPS, 1 uM PMSF, 1 mM Biotin, Protease Inhibitor Cocktail). Bacteria were lysed by sonication (4 x 3 min, 50% amplitude, 01 s on 01 s off). Proteins were purified by Ni-NTA (2 x HiTrap Chelating HP, 5 mL, GE Healthcare), buffer exchanged to cleavage buffer (20 mM TrisHCl pH 7.4, 100 mM NaCl, 1 mM TCEP), and cleaved with an in-house 3C protease (H10-GST-3C) overnight at 4°C. The cleaved protein was again loaded to a Ni-NTA to bind the cleaved H6-MBP and the H10-GST-3C. The flow-through containing mSA2 was collected. The mSA2-containing fractions were pooled, concentrated and loaded onto a gel filtration column (HiLoad 16/600 Superdex 75 pg) equilibrated with phosphate buffered saline (PBS) buffer pH 7.4 (containing 0.5 mM TCEP). The mSA2-containing fractions were concentrated in an Amicon Ultra-15 centrifugal filter device (3 kDa MW cutoff) and either directly measured by NMR or buffer exchanged (PBS^100%^*^D^*^2^*^O^*) using the Amicon Ultra-15 centrifugal filter device. Protein concentration was measured using a NanoDrop. For apo-mSA2, no biotin was added to the expression medium, and the sample was used immediately after the second Ni-NTA column after the cleavage of the H6-MBP tags due to the limited stability of apo-mSA2 in solution.

### NMR Spectra Acquisitions

1D and NOESY experiments were carried out at 298 K on a Bruker 600 MHz spectrometer equipped with a TCI cryoprobe. The initial sample concentration was 860 *µ*M. For 1D experiments, the number of scans was set to 512. For the NOESY experiment, the number of scans was set to 32, the size of fid was set to 2048, 512 (F2, F1) and the mixing time was set to 100 ms.

### Ligand ^1^H-NMR chemical shifts computation

The X-ray crystal structure of biotin bound to the mutant streptavidin mSA2 (PDB ID: 4JNJ)^22^ was used as the starting model for the computational study. The protein was protonated at pH 7.4 using the H++ web server (version 4.0),^61^ while the ligand was protonated using the ProteinsPlus server.^62–65^ The restrained electrostatic potential (RESP) point charges for biotin were computed by means of HF/6-31G(d) on the optimized geometry of the ligand at the B3LYP-D3/6-311G(d,p) level of theory using Gaussian16. ^66^ The Amber ff19SB force field^67^ was applied to the protein, while the General Amber Force Field (GAFF)^68^ was used for the ligand. The system was neutralized by adding chloride ions and solvated in an octahedral box of TIP3P water molecules, ^69^ extending 10 Å from the solute, using the LEaP module of AmberTools21.^70^ Water molecules present in the crystal structure were retained. Energy minimization was conducted under periodic boundary conditions using the Amber20 software.^70^ Non-bonded interactions were calculated with a 10 Å cutoff, while long-range electrostatic interactions were treated using the smooth Particle Mesh Ewald (PME) method.^71^ The SHAKE algorithm was applied to constrain all bonds involving hydrogen atoms.^72^ Minimization was performed in two stages: initially, 10,000 cycles using the steepest descent algorithm, followed by 10,000 cycles with the conjugate gradient method. In the first stage, only hydrogen atoms were minimized, followed by the minimization of solvent molecules in the second stage.

The ligand, accompanied by two water molecules H-bonded to its carboxylate group, was chosen for further QM calculations, alongside interacting amino acids: Asn45, Ala47, Trp79, Cys86, Ser88, Thr90, Trp108, Leu110, Asn23, Ser27, Tyr43, Trp92, Asp128, Thr48, Gly49 and Cys50. The side chains up to the C*_α_* atoms of Asn45, Ala47, Trp79, Cys86, Ser88, Thr90, Trp108, and Leu110 were included, with the C*_α_*-N and C*_α_*-C*_Carbonyl_* bonds being cleaved and filled with hydrogen atoms. For Asn23, Ser27, Tyr43, Trp92, and Asp128, the bonds between C*_β_* and C*_α_* were cut and filled with hydrogen atoms. Additionally, the sequence Thr48-Gly49-Cys50 underwent cleavage at the C*_α_*-N bond of Thr48 and the C*_α_*- C*_Carbonyl_* bond of Cys50, followed by filling with hydrogen atoms.

The isotropic chemical shielding values were determined using the Gauge-Independent Atomic Orbital (GIAO) approach^53–57^ and the wB97XD functional^58^ in combination with the def2-TZVP basis set, as implemented in Gaussian16. ^66^ The choice of theory level is based on the benchmark carried out in our previous study.^39^

To compare the prediction with experimental data, the calculated ^1^H NMR chemical shielding values (*σ*) need to be converted to chemical shifts (*δ*) using a standard reference:

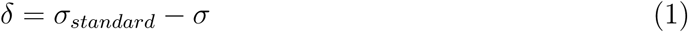

In this study, we used a linear model with a fixed slope of -1 to calculate the chemical shielding of the reference standard, specifically water.^39^

### ^1^H-NMR chemical shifts decomposition

Building on the approach outlined by Scheiner,^60^ we decompose the chemical shift of protons in the bound form into two physically meaningful components: the “Environmental shielding effect” and the “Deshielding due to H-bond formation,” as referenced further in the text. Following the setup of the QM region, we computed the “Environmental shielding effect” by removing the ligand and replacing its C-bound protons with “dummy atoms” positioned at the original proton locations. Shielding was then calculated at these precise positions. This effect arises from the electron density of hydrogen-accepting groups (here, interacting residues), which can respond to external magnetic fields even without forming a hydrogen bond (HB) or having a proton donor nearby. Therefore, this positional effect is independent of any hydrogen bond formation.

If a hydrogen bond does form, there are additional contributions from charge transfer between the donor and acceptor, along with internal polarization changes within each group. This "Deshielding due to H-bond formation" was calculated by subtracting two components from the chemical shifts computed for the entire ligand-binding pocket system: (1) the chemical shifts of the ligand alone and (2) “Environmental shielding effect”. As demonstrated in Scheiner’s work,^60^ this component serves as an indicator of the presence and strength of a hydrogen bond, as it has shown correlation with binding energy in model systems.

### Extended Transition State Natural Orbitals for Chemical Valence (ETS-NOCV) charge and energy decomposition method

Extended Transition State (ETS)^73^ is applied to partition the total binding energy between interacting fragments into chemically meaningful contributions:

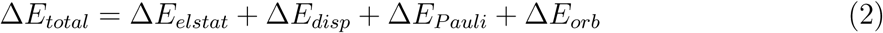

The first term, Δ*E_elstat_*, in the equation represents the classical electrostatic interactions between selected fragments. The Δ*E_disp_* concerns semiempirical Grimme dispersion correction (D3).^74^ The third component, Δ*E_Pauli_*, accounts for Pauli repulsion between occupied orbitals of fragments. The last, orbital interactions contribution, Δ*E_orb_*, is stabilizing and covers the interaction energy between occupied orbitals of the first fragment with unoccupied orbitals of the second one and vice versa, as well as the polarization effects. Natural Orbitals for Chemical Valence (NOCV) theory^75^ enables the decomposition of the overall deformation density, Δ*ρ_orb_* = *ρ* − *ρ*_0_ (where *ρ* is a density of the molecule and *ρ*_0_ is a density of non-interacting fragments), into the distinct bonding channels such as *σ*, *π*, *δ*, etc: Δ*ρ_orb_* = ∑*_i_* Δ*ρ_orb_*(*i*).

Finally, combining the ETS method with Natural Orbitals for Chemical Valence (NOCV) theory,^40^ allows us to quantify the energy values of the Δ*E_orb_*, corresponding to Δ*ρ_orb_*(*i*) contributors: Δ*E_orb_* = ∑*_i_* Δ*E_orb_*(*i*). Overall, the ETS-NOCV method provides a qualitative and quantitative, picture of a chemical bonding.

ETS-NOCV analyses were performed using the Amsterdam Density Functional (ADF) software, version 2023.104. ^76,77^ The BLYP functional, coupled with Grimme’s D3 dispersion correction with the Becke-Johnson damping, ^74^ was applied for bonding analysis due to its recognized reliability in describing non-covalent interactions.^78,79^ All calculations utilized a triple-*ζ* STO basis set with polarization functions (TZP), and were conducted without imposing symmetry constraints, using an all-electron basis set.

### The Domain-based Localized Pair-Natural Orbital Singles and Doubles Coupled Cluster with perturbative Triples (DLPNO-CCSD(T)) and Local Energy Decomposition

DLPNO-CCSD(T)^41–45^ Local Energy Decomposition (LED)^46–48^ calculations were performed in ORCA 5.0.4 program,^80^ using def2-TZVP basis,^81,82^ RIJCOSX approximation^83^ and TightPNO settings for PNO convergence.

The Domain-based Localized Pair-Natural Orbital Singles and Doubles Coupled Cluster with perturbative Triples (DLPNO-CCSD(T)) is a local approximation of the Coupled Clusters method, which itself is often referred to as a ‘golden standard’ for quantum-chemical calculations. Its main feature is the localization of the orbitals and generation of the Projected Atomic Orbitals (PAOs) from occupied localized orbitals, followed by creation of Pair Natural Orbitals from selected PAOs.

Local Energy Decomposition^46–48^ is defined within the DLPNO-framework, and decomposes the interaction energy into the Hartree-Fock and correlation energies:

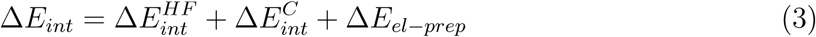

Electronic preparation terms of the distorted fragments 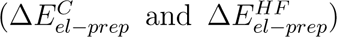 were added for clarity. Hartree-Fock interaction energy consists of the electrostatic and exchange terms:

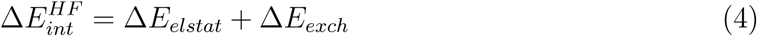

whereas the correlation interaction consists of the following terms:

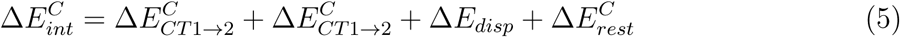

where 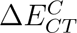 terms correspond to the instantaneous strong-pair charge transfer terms, Δ*E_disp_* is a sum of dispersion stemming from strong and weak pairs, and 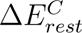 is a summation of the non-dispersive part of weak-pair interaction energy and the triples correction.

## Acknowledgement

The authors would like to thank Mariusz P. Mitoraj and Moriz Mayer for their scientific input and critical revision of the manuscript. A. L. Ptaszek was funded by the Christian Doppler Laboratory for High-Content Structural Biology and Biotechnology, Austria. A. L. Ptaszek and Pedro A. Sánchez-Murcia thank the Medical University of Graz for computation time at the MedBioNode cluster. Parts of the calculations were carried out with the equipment purchased thanks to the financial support of the European Regional Development Fund in the framework of the Polish Innovation Economy Operational Program (contract no. POIG.02.01.00-12-023/08). We gratefully acknowledge Polish high-performance computing infrastructure PLGrid (HPC Centers: ACK Cyfronet AGH) for providing computer facilities and support within computational grant no. PLG/2024/016976.

## Supporting Information

Contents include details of mSA2 mutant similarity to WT streptavidin, NMR spectra and measured chemical shifts, LED/DLPNO-CCSD(T) results, DID interfragment dispersion visualisation, more detailed NOCV analysis and CSP predictions using the Pople model.

CSs: chemical shifts
CSP: chemical shift perturbations
NMR: Nuclear Magnetic Resonance
WT: wild-type

