## Supporting Information for "From Weak Interactions to Strong Affinity: Deciphering the Streptavidin-Biotin Interaction through NMR and Computational Analysis"

⊥*BioTechMed-Graz, Mozartgasse 12/II, A-8010 Graz, Austria*

#*Equal contribution*

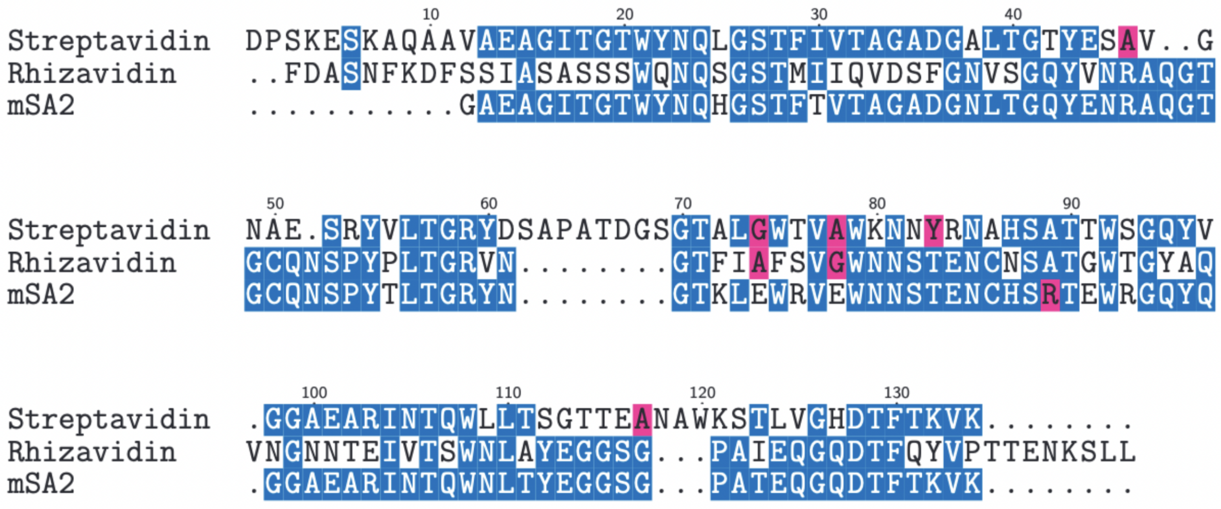


Figure S1: Sequence similarities of WT streptavidin, WT rhizavidin and mSA2.[S1]

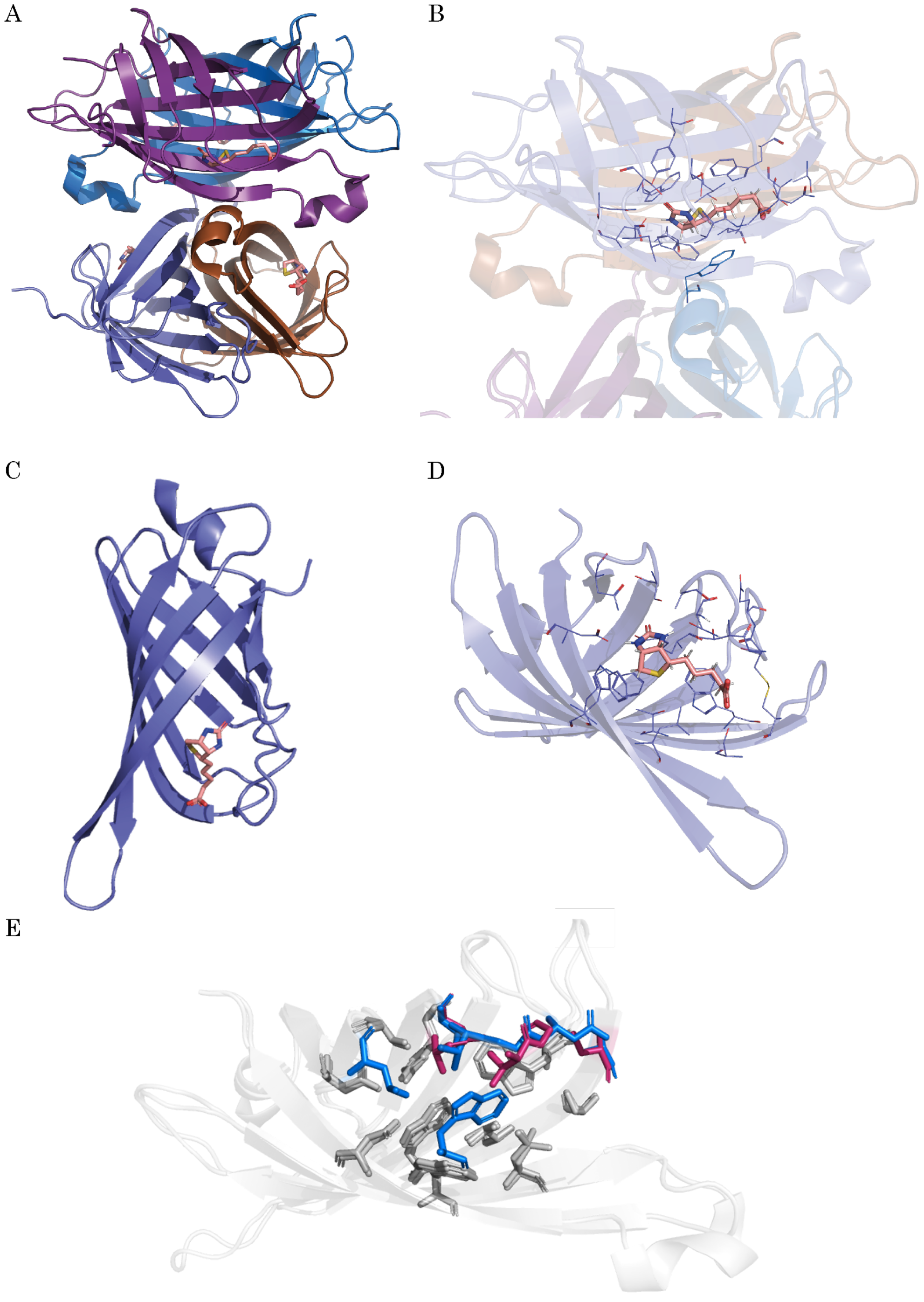


Figure S2: (A) The crystal structure of WT streptavidin is shown as a homotetrameric protein with (B) the binding pocket of biotin bound to the WT streptavidin. (PDB ID: 3RY2) (C) The crystal structure of the mSA2 mutant is shown, with (D) the binding pocket of biotin. (PDB ID: 4JNJ) (E) The aligned structures of the WT and mutant are illustrated, where the differences in amino acids in the biotin-binding pocket are highlighted in blue for the WT and magenta for the mSA2 mutant.

### HSQC

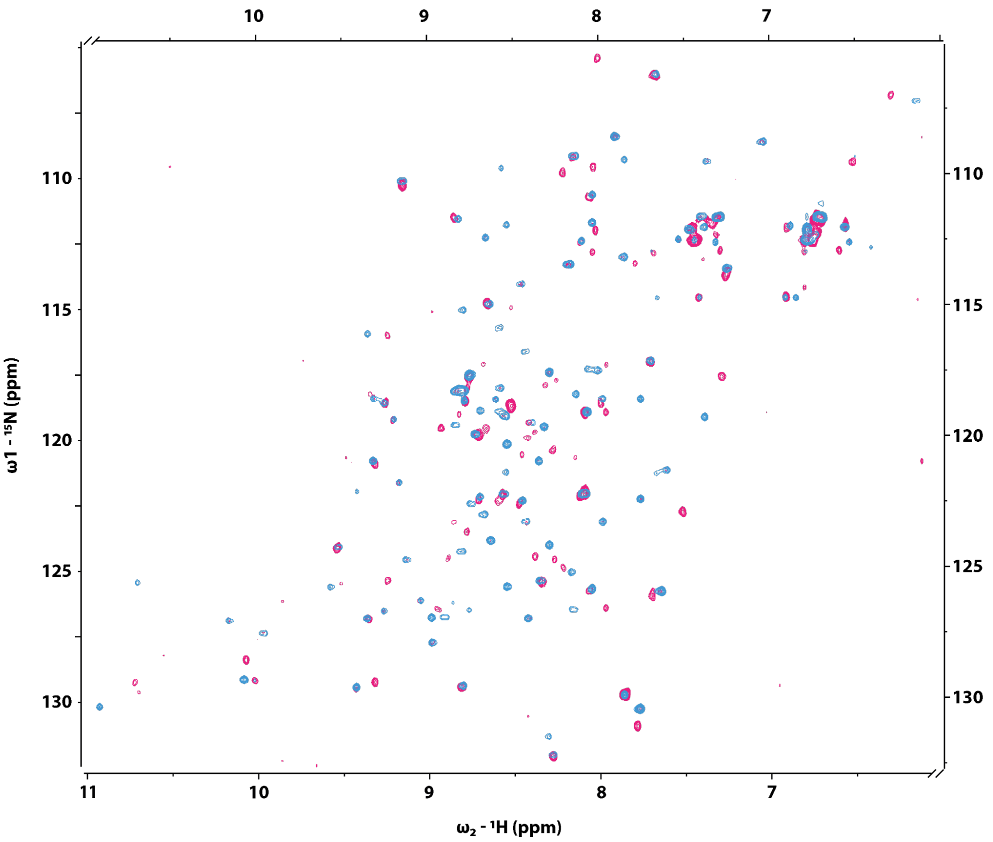


Figure S3: $^{1}$H-$^{15}$N HSQC of mSA2-apo (pink) and mSA2-biotin (blue).

### NOESY

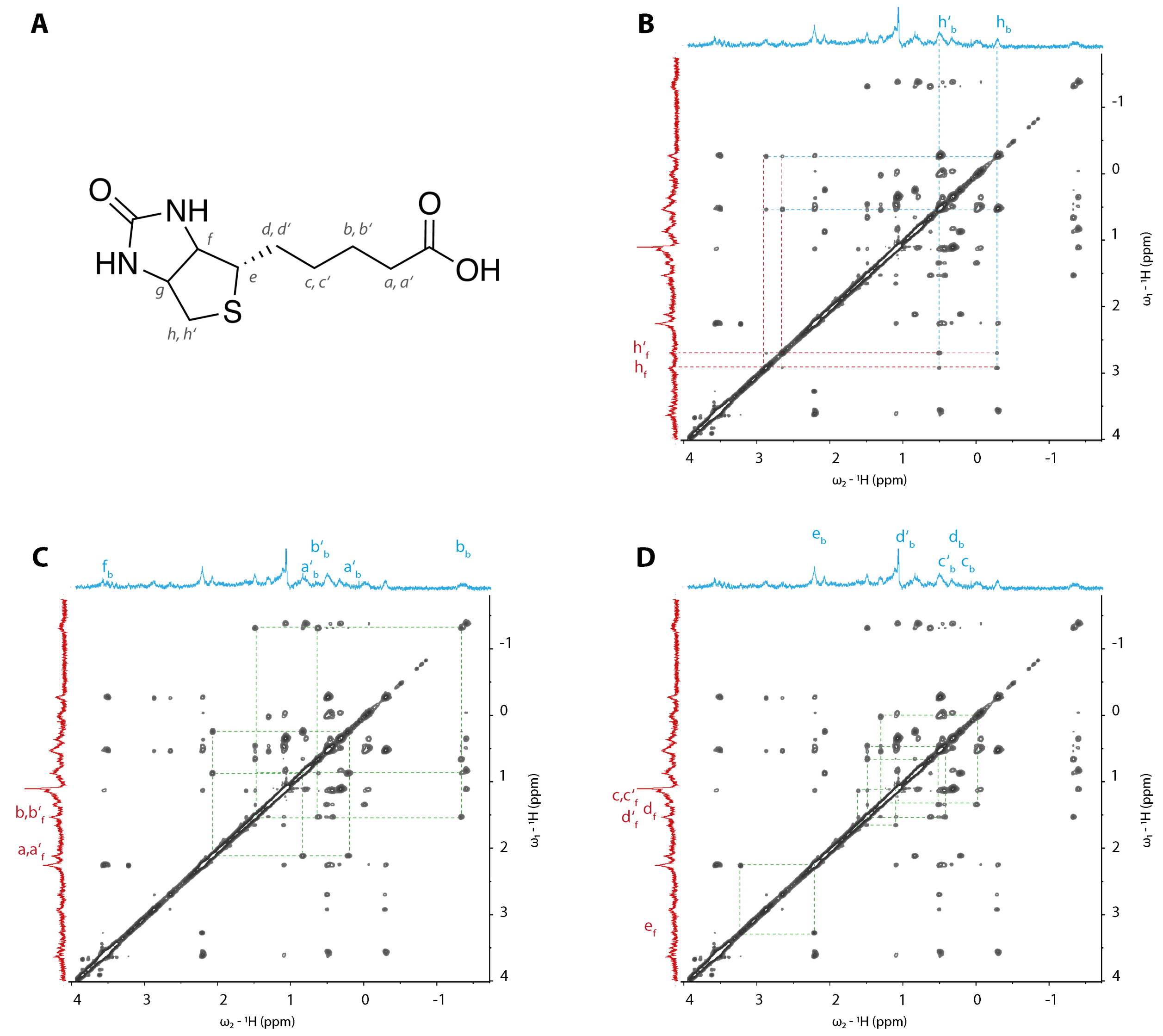


Figure S4: $^{1}$H-$^{1}$H NOESY spectra of biotin in complex with mSA2. Spectra were recorded as mSA2-biotin 1:1 complex, due to partial precipitation of mSA2, biotin is in 1.5-fold excess. (A) Chemical structure of biotin: protons are numbered with letters a - h'. (B) Scheme for the assignment strategy of NOE cross peaks showing exchange peaks between the free signals (red) and the bound signals (blue), NOE between two bound signals, and NOE between free to relay bound signals. (C and D) CSPs between free and bound signals are shown in dotted green lines.

Table S1: Experimentally determined CS and CSP for biotin's carbon-bound H-atoms.

| **Proton** | $\delta$ Biotin (ppm) | $CSP_{Experimental}$ (ppm) | $\delta$ Biotin:mSA2 (ppm) |
|---|---|---|---|
| a | 2.13 | -1.88 | 0.25 |
| a´ | 2.13 | -1.28 | 0.85 |
| b | 1.49 | -2.89 | -1.41 |
| b´ | 1.49 | -0.82 | 0.67 |
| c | 1.30 | -1.31 | -0.01 |
| c´ | 1.30 | -0.82 | 0.48 |
| d | 1.52 | -1.17 | 0.35 |
| d´ | 1.61 | -0.49 | 1.12 |
| e | 3.25 | -0.98 | 2.27 |
| f | 4.35 | -0.70 | 3.65 |
| g | 4.50 | -0.94 | 3.56 |
| h | 2.89 | -3.17 | -0.28 |
| h´ | 2.66 | -2.15 | 0.51 |

Table S2: LED/DLPNO-CCSD(T) energy decomposition describing the interaction between the fragments analogous to ones described in main manuscript: biotin together with nearby water molecules (fragment 1 in all fragmentations) and the binding pocket (fragment 2); biotin with waters and the ureido subpocket (Asn23, Ser27, Tyr43, Asn45, Trp92, Asp12); biotin with waters and the tail subpocket (Ala47, Thr48, Gly49, Cys50, Cys86, Trp79, Ser88, Thr90, Trp108, Leu110). Energies listed in kcal/mol.

| **Energy term** | **Biotin+$H_2O$ $\cdots$ pocket** | **Biotin+$H_2O$ $\cdots$ ureido subpocket** | **Biotin+$H_2O$ $\cdots$ tail subpocket** |
|---|---|---|---|
| $\Delta E_{int}$ | -83.23 | -22.18 | -58.71 |
| $\Delta E_{elst}^{HF}$ | -348.78 | -170.09 | -182.48 |
| $\Delta E_{exch}^{HF}$ | -71.28 | -31.95 | -39.95 |
| $\Delta E_{disp}$ | -62.53 | -20.14 | -42.82 |
| $\Delta E_{CT1\rightarrow 2}^{C}$ | -41.42 | -16.74 | -24.71 |
| $\Delta E_{CT2\rightarrow 1}^{C}$ | -31.85 | -16.30 | -14.83 |
| $\Delta E_{rest}^{C}$ | -38.13 | -12.62 | -21.76 |
| $\sum \Delta E_{el-prep}$ | 510.76 | 245.67 | 268.85 |

### LED analysis

DLPNO-CCSD(T)-based Local Energy Decomposition is a powerful tool that allows for an unprecedented level of detail in describing London dispersion.[S2–S9] Although not strictly additive, the contributions from biotin-ureido subpocket and biotin-tail subpocket models sum up almost perfectly to their counterparts in the whole model, allowing us to draw conclusions on their respective magnitudes, Table S2. Immediately noticeable is the greater stabilization stemming from the tail part of binding site. Almost all energy terms are more stabilizing for the tail part, owing to greater number of contacts in this region, showing clearly, that many weaker interactions can dominate over the regular hydrogen bonds. Particularly, the dispersion energy is twice as high as for the biotin-ureido subpocket, followed by significantly higher exchange energy term, jointly indicating non-classical nature of the said stabilization. Another interesting difference between the fragmentations is the prevalence of biotin→binding site correlation charge-transfer term (which can be interpreted as the instantaneous ion pair formation), which is consistent with the direction of typical charge-transfer.[S9] It may be due to the negative charge on biotin, which probably increases charge-transfer in this direction.

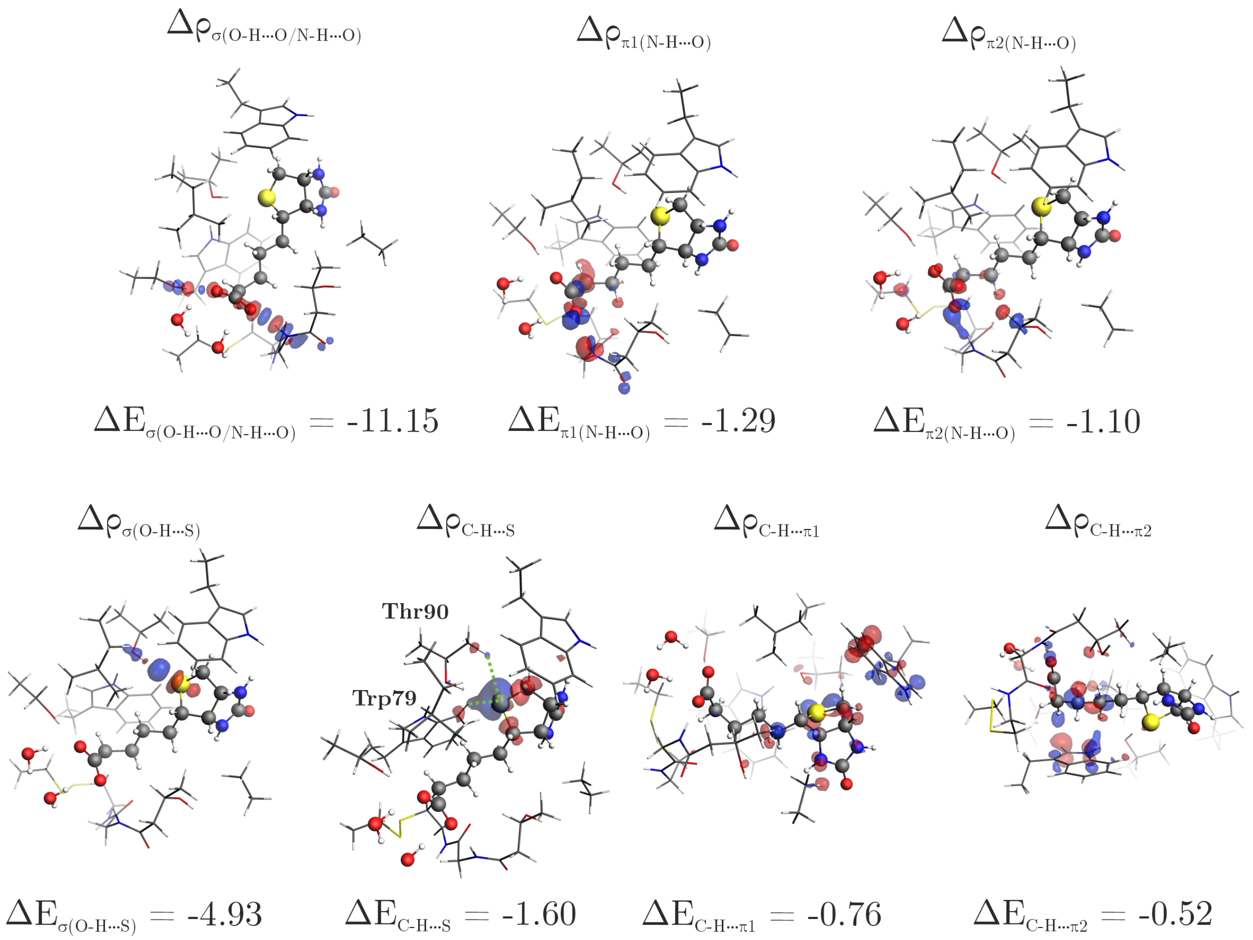


Figure S5: Selected NOCV deformation density channels along with the energy contributions of biotin molecule and two water molecules (fragment 1) interacting with the protein tail subpocket (fragment 2). The biotin and water molecules (fragment 1) are represented by balls and sticks, and the residues from the binding site (fragment 2) are represented by sticks.

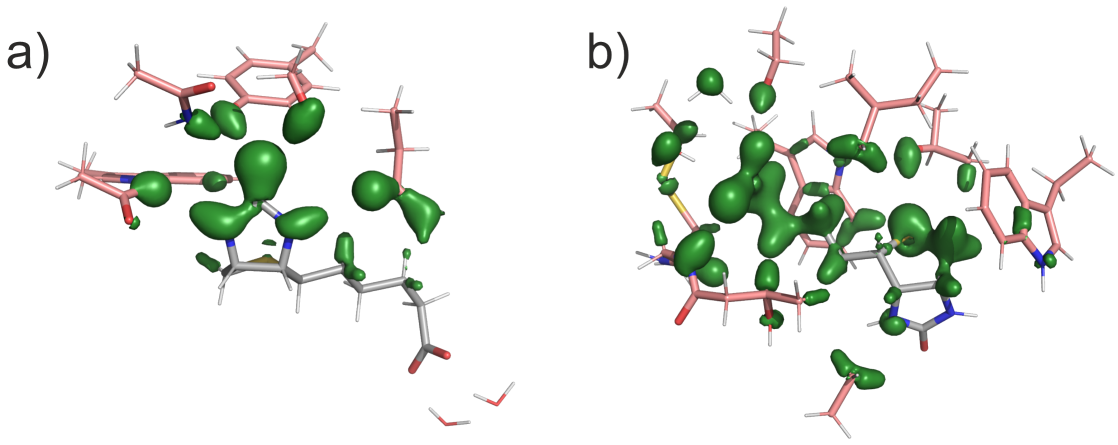


Figure S6: DID interfragment dispersion visualization of biotin and water molecules interaction with a) the ureido subpocket and b) the tail subpocket. The biotin and water molecules (fragment 1) are coloured gray, and the residues from the binding site (fragment 2) are coloured pink. LED-DLPNO-CCSD(T)/def2-tzvp.

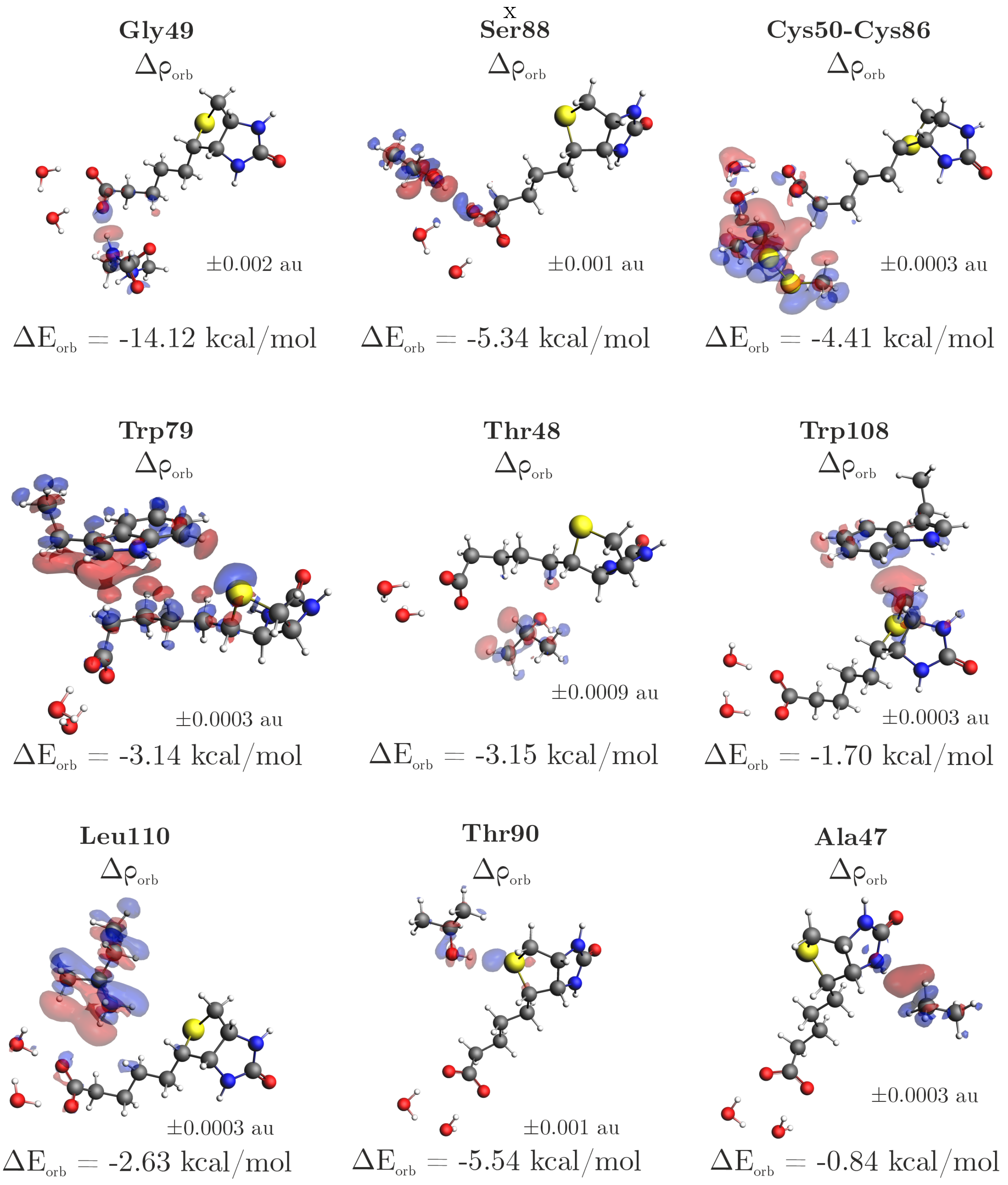


Figure S7: The overall deformation density $\Delta\rho_{orb}$ with the corresponding $\Delta E_{orb}$ of biotin molecule and two water molecules (fragment 1) interacting with individual residues from the tail subpocket (fragment 2).

**Pople's Model**

In Pople's equation, the center of the aromatic ring induces a magnetic field according to Equation 1:[S10]

$$\Delta\sigma = 10^6 \times \frac{ne^2a^2}{4\pi mc^2} \times \frac{3cos^2\theta - 1}{r^3} \quad (1)$$

In Equation 1, $\Delta\sigma$ represents the change in the chemical shift in ppm, $n$ equals the number of $\pi$-electrons (6 for each of the aromatic rings of Trp), $e$ equals the elementary charge in Franklin (1 e = 4.803$\times$ $10^{10}$ Franklin), $a$ equals the radius of the aromatic rings in Trp (6-ring: 1.39$\times10^{-8}$ cm; 5-ring: 1.22$\times10^{-8}$ cm), $m$ equals the electron mass in gram (9.109$\times10^{-28}$ g), $c$ is the speed of light (2.998$\times10^{10}$ m$\times$s$^{-1}$), $\theta$ represents the angle between the normal vector of the plane of one of the aromatic rings of Trp and the proton to ring center vector in rad, and $r$ represents the distance from the proton to either the 5-ring or 6-ring center of the aromatic ring. Values (r, $\theta$ and a) were extracted from the crystal structure (PDB ID: 4JNJ).

Table S3: CSP prediction from the Pople model.

| Proton | $CSP_{Experimental}$ (ppm) | $CSP_{Pople}$ (ppm) |
|---|---|---|
| a | -1.88 | -2.40 |
| a´ | -1.28 | -0.84 |
| b | -2.89 | -2.12 |
| b´ | -0.82 | -0.70 |
| c | -1.31 | -0.86 |
| c´ | -0.82 | -0.48 |
| d | -1.17 | -0.51 |
| d´ | -0.49 | -0.40 |
| e | -0.98 | -0.56 |
| f | -0.70 | -0.27 |
| g | -0.94 | -0.64 |
| h | -3.17 | -3.94 |
| h´ | -2.15 | -1.86 |

(S2) Neese, F.; Wennmohs, F.; Hansen, A. Efficient and accurate local approximations to coupled-electron pair approaches: An attempt to revive the pair natural orbital method. *The Journal of Chemical Physics 130*, 114108.

(S3) Neese, F.; Hansen, A.; Liakos, D. G. Efficient and accurate approximations to the local coupled cluster singles doubles method using a truncated pair natural orbital basis. *The Journal of Chemical Physics 131*, 064103.

(S4) Riplinger, C.; Sandhoefer, B.; Hansen, A.; Neese, F. Natural triple excitations in local coupled cluster calculations with pair natural orbitals. *The Journal of Chemical Physics 139*, 134101.

(S5) Guo, Y.; Riplinger, C.; Becker, U.; Liakos, D. G.; Minenkov, Y.; Cavallo, L.; Neese, F. Communication: An improved linear scaling perturbative triples correction for the domain based local pair-natural orbital based singles and doubles coupled cluster method [DLPNO-CCSD(T)]. *The Journal of Chemical Physics 148*, 011101.

(S6) Saebo, S.; Pulay, P. Local Treatment of Electron Correlation. *Annual Review of Physical Chemistry 44*, 213–236.

(S7) Altun, A.; Saitow, M.; Neese, F.; Bistoni, G. Local Energy Decomposition of Open-Shell Molecular Systems in the Domain-Based Local Pair Natural Orbital Coupled Cluster Framework. *Journal of Chemical Theory and Computation 15*, 1616–1632.

(S8) Altun, A.; Izsák, R.; Bistoni, G. Local energy decomposition of coupled-cluster interaction energies: Interpretation, benchmarks, and comparison with symmetry-adapted perturbation theory. *International Journal of Quantum Chemistry 121*, e26339.

(S9) Schneider, W. B.; Bistoni, G.; Sparta, M.; Saitow, M.; Riplinger, C.; Auer, A. A.; Neese, F. Decomposition of Intermolecular Interaction Energies within the Local Pair Natural Orbital Coupled Cluster Framework. *Journal of Chemical Theory and Computation 12*, 4778–4792.

(S10) Pople, J. Proton magnetic resonance of hydrocarbons. *The Journal of Chemical Physics* **1956**, *24*, 1111–1111.
